# ssGSEA3.0: Feature-Level Bootstrap Resampling for Robust Assessment of Statistical Significance

**DOI:** 10.64898/2026.10.28.762524

**Authors:** Natalie M. Clark, D. R. Mani

## Abstract

**Motivation:** Single-sample Gene/Post-Translational Modification Set Enrichment Analysis (ssGSEA/PTM-SEA) methods assess pathway or signature activity per sample using permutation-based significance testing. ssGSEA2.0’s weighted scoring (ssGSEA2.0-orig) uses a correlation-vector reordering that inflates false positives at nonzero weight; directly fixing this (ssGSEA2.0-fixed) restores null-distribution calibration but sacrifices detection power at nonzero weight.

**Results:** ssGSEA3.0 replaces feature-label (gene- or site-level) permutation with feature-level bootstrap resampling, which remains calibrated across weighting schemes. On published lung adenocarcinoma and phosphoproteomic perturbation datasets, ssGSEA3.0 shows greater detection power than ssGSEA2.0-fixed, with an additional gain for directional (PTM-SEA) scoring. Previously published findings remain robust across methods, including at nonzero weight. Bootstrapping adds a ∼1.5- to 2-fold computational cost.

**Availability and Implementation:** ssGSEA3.0 is implemented as an open-source R package (github.com/broadinstitute/ssGSEA3.0, DOI: 10.5281/zenodo.23734262) and includes the worked examples on biological data presented here.

## Introduction

Gene set enrichment analysis (GSEA) is extensively used to determine pathway enrichment in phenotype classes of interest, providing a more systems-level view of differential gene expression (Subramanian *et al*. 2005). Single-sample GSEA (ssGSEA) adapted this framework to score gene set or pathway activity within a single sample rather than across a two-class comparison, yielding one enrichment score per sample per gene set (Barbie *et al*. 2009). ssGSEA2.0 introduced permutation-based significance testing for these enrichment scores and extended the approach to site-centric phosphoproteomic data through PTM-SEA, which incorporates the directionality of a post-translational modification by combining separately scored “up” and “down” pathway members (Krug *et al*. 2019). These implementations compute an enrichment score in which each feature’s contribution is scaled by a weight parameter, with nonzero weight favoring features with larger expression magnitude and weight = 0 reducing to a rank-only score. Significance is assessed against a null distribution derived by permuting gene or site labels.

The permutation implementation in ssGSEA2.0 (ssGSEA2.0-orig) shuffles each feature’s expression-based weight along with its rank. For weight > 0, this permutation scheme causes highly expressed features to drive the permuted score, resulting in a null distribution that is too narrow and inflates false positives (GitHub issue #45; github.com/broadinstitute/ssGSEA2.0/issues/45). Because nonzero weighting is the default recommended for continuously valued expression or abundance data, this miscalibration potentially affects previously published ssGSEA and PTM-SEA analyses. A straightforward fix (ssGSEA2.0-fixed) is to retain the expression values in their original order and permute only the ranking of the genes or sites. Thus, gene/site order and expression magnitude (weight) are decoupled during permutation. While this restores calibration at nonzero weight and reins in false positives, as shown below, the method suffers from reduced power to detect true pathway enrichments.

We show here that feature-level bootstrap resampling provides an effective alternative to permutation testing, and introduce ssGSEA3.0, an open-source R package (github.com/broadinstitute/ssGSEA3.0) that replaces gene-label permutation with feature-level bootstrap resampling for both non-directional (ssGSEA) and directional (PTM-SEA) single-sample enrichment scoring. Using synthetic simulations and two previously published biological datasets, lung adenocarcinoma proteogenomic profiles (Satpathy *et al*. 2025) and a PI3Kα-inhibitor phosphoproteomics dataset (Krug *et al*. 2019), we show that ssGSEA3.0 remains calibrated while showing greater detection power than the corrected permutation procedure across weighting schemes, with an additional power gain specific to directional PTM-SEA scoring. We further characterize the computational cost of bootstrap resampling relative to permutation, providing ssGSEA3.0 as a reproducible, drop-in replacement for existing ssGSEA2.0 workflows.

## Methods

### ssGSEA implementations

We compare three implementations of single-sample enrichment scoring using the following naming convention throughout: ssGSEA2.0-orig (ssGSEA2.0 as published at github.com/broadinstitute/ssGSEA2.0 and in (Krug *et al*. 2019)), ssGSEA2.0-fixed (a branch of ssGSEA2.0 with the correlation-vector reordering fixed; 10.5281/zenodo.23734330), and ssGSEA3.0 (our new implementation). All three compute a weighted running-sum enrichment score (ES) in which each feature’s contribution is scaled by a weight parameter (with weight = 0 denoting unweighted rank-based scoring). Significance is assessed against a null distribution built either by gene-label permutation (ssGSEA2.0) or by feature-level bootstrap resampling (ssGSEA3.0). The permutation procedure in ssGSEA2.0-orig reorders the per-feature vector of magnitudes used to weight each feature’s contribution to the running-sum enrichment score (referred to throughout this work as the correlation-vector) together with the permuted feature ranks. ssGSEA2.0-fixed corrects this by holding the weight vector fixed to its observed order and permuting only gene-set membership.

### Feature-level bootstrap resampling

ssGSEA3.0 replaces gene-label permutation with bootstrap resampling of the measured features themselves (genes for ssGSEA, PTM sites for PTM-SEA) sampled with replacement from the full measured set. For each of *n* replicates, a bootstrap draw is taken, re-ranked by its resampled expression values, and the gene set’s members are relocated within that ranking (for directional scoring, the up- and down-regulated portions are rescored separately from the same draw). Because each feature’s weight is always associated with its original value, in its original order, rather than reshuffled along with the resampled ranks, this procedure avoids the artifacts arising from gene-label permutation.

The bootstrap *p*-value calculation uses the shifted null approach, where the *n* bootstrap enrichment scores are recentered to their mean thereby approximating a permutation-style null distribution centered at zero (Efron and Tibshirani 1994). Because a naive percentile-based bootstrap p-value is known to be under-dispersed and overly significant, we instead use the bootstrap standard error as the centered distribution’s standard deviation and compute a *z*-score-based *p*-value from the normal distribution (Efron 1982; MacKinnon 2002). Finally the normalized enrichment score (NES) for the observed enrichment statistic is obtained by dividing the score by the mean of the same-signed centered bootstrap scores.

### Analysis parameters

Unless otherwise noted, all analyses employ representative parameters commonly used for most analyses: *nperm* = 1000 (bootstrap replicates for ssGSEA3.0; permutations for ssGSEA2.0), *sample.norm.type* = rank (each sample’s feature values are rescaled to within-sample ranks, which then serve as the weighting values when weight > 0), and *statistic* = area.under.RES (running enrichment score test statistic). *correl.type* = z.score for non-directional (ssGSEA) scoring and rank for directional (PTM-SEA) scoring (governs how the per-feature weight vector is built when weight > 0), matching each mode’s convention. *min.overlap* (the minimum required overlap between a gene set and the measured features) is 10 for the LUAD analyses (ssGSEA) and 5 for the PI3K analysis (PTM-SEA). ssGSEA3.0 additionally uses *seed* = 42 for reproducibility; ssGSEA2.0 has no *seed* parameter.

### Power evaluation using biological datasets

We used two previously published datasets to evaluate the agreement of the three implementations as well as their ability to recapitulate known biology. The first dataset contains the feature weights from non-negative matrix factorization (NMF) of a lung adenocarcinoma (LUAD) cohort (Satpathy *et al*. 2025) derived using the PANOPLY pipeline (Mani *et al*. 2021). The original ssGSEA analysis presented in the paper used weight = 1 and the MSigDB (Liberzon *et al*. 2015) HALLMARK collection. In addition to repeating the analysis on the HALLMARK collection (50 curated, similarly sized gene sets), we performed a second analysis using the same weight parameter on the MSigDB REACTOME collection (1,167 gene sets ranging from ten to several hundred genes). The second dataset is a site-centric phosphoproteomics dataset from the original PTM-SEA publication (Krug *et al*. 2019) in which T47D breast cancer cells were treated with the PI3Kα inhibitor BYL719 versus DMSO control at 6h and 24h. As the input values of this dataset are log-signed *p-*values from the moderated two-sample *t*-test at each timepoint (which are generally not used as weights since they are not measured values), we performed PTM-SEA with weight = 0 against PTMsigDB v2.0.0’s human flanking-sequence signatures (<u>proteogenomics.shinyapps.io/ptmsigdb/</u>).

Power was assessed by determining the number of gene sets significant at a dataset-specific FDR threshold (LUAD HALLMARK: FDR < 0.01, matching the source publication’s own threshold; LUAD REACTOME and PI3K PTMsigDB: FDR < 0.05). Agreement between implementations was quantified by Pearson and Spearman correlation of NES as well as the overlap between significant gene sets at the relevant FDR threshold.

### Calibration and power evaluation using synthetic simulations

Two synthetic simulations were performed to further evaluate the three implementations. The first simulation was used to determine each implementation’s calibration against null gene sets. Four hundred gene sets were randomly constructed based on the genes detected in the LUAD dataset with sizes log-uniform between 10-300 features to mimic REACTOME’s own size diversity. The LUAD NMF feature weights were then scored against these randomly constructed gene sets at weight = 0 and 0.75, with weight = 0 serving as a negative control at which the correlation-vector reordering does not manifest. Calibration was assessed two ways: the nominal rate (the fraction of null gene sets called significant at *p* < 0.05, expected ∼5%) and the Kolmogorov-Smirnov test, which compares the entire null *p*-value distribution against Uniform(0,1).

To determine whether ssGSEA3.0’s power advantage over ssGSEA2.0-fixed reflects a genuine improvement in detection power rather than bootstrap miscalibration, we used a synthetic ground-truth simulation. A fully synthetic dataset was generated: single gene sets were defined by drawing a background of 2,000 features from an *i.i.d*. Normal(0,1) distribution and then randomly selecting 20 or 100 of those 2,000 features. We first examined differences in detection power using non-directional gene sets, which are used for ssGSEA. We embedded a ± 0.5 shift in gene set members in each of 10,000 gene sets (20 members per gene set, *nperm* = 1000) where the shift sign was randomized per gene set and applied uniformly to all members of each trial’s gene set. We then performed ssGSEA using weight = 1, and compared the methods using a paired McNemar-test. For directional gene sets used in PTM-SEA, to address potential miscalibration, we first performed a null-calibration simulation (500 randomly generated gene sets, *nperm* = 1000) at two gene set sizes (20 and 100 members) for both directional and non-directional gene sets with both permutation (ssGSEA2.0-fixed) and bootstrap (ssGSEA3.0) approaches. Then, we used the same shifted gene set approach as before but added a directional component where the shift sign was fixed by membership (positive for “up” and negative for “down” members) and used weight = 0. Bootstrap versus permutation detection power was compared using a paired McNemar test for directional and non-directional scoring separately.

### Computational timing

Wall-clock timing was measured on a single machine (Apple M1 Max MacBook Pro, 10 cores [8 performance + 2 efficiency], 32GB memory, macOS 15.7.7, R 4.4.2). Both ssGSEA2.0-orig and ssGSEA3.0 were run with parallelization enabled across all 10 cores (*par* = TRUE, *spare.cores* = 0). ssGSEA2.0-fixed was not separately timed as it differs from ssGSEA2.0-orig only in holding the weight vector fixed, which does not significantly affect the runtime. Each dataset was timed end-to-end via explicit wall-clock timestamps. Timings reflect single runs on one machine rather than averages over repeated trials, and should be read as representative rather than as precision benchmarks.

### Implementation and reproducibility

ssGSEA3.0 is implemented as an open-source R package (github.com/broadinstitute/ssGSEA3.0, DOI: 10.5281/zenodo.23734262). Gene sets are scored in parallel across available CPU cores or serially and, when a random seed is supplied, produce identical output regardless of the number of cores used or how work is scheduled across them. Within the bootstrap replicate loop, each feature’s position is resolved via a hashed lookup against the gene set’s unique members rather than a linear scan of all measured features, reducing computational time without altering output.

## Results

### The permutation approach in ssGSEA2.0-orig is miscalibrated for a nonzero weight

The current permutation approach implemented in the published ssGSEA2.0 algorithm (referred to throughout this manuscript as ssGSEA2.0-orig) introduces a correlation-vector reordering that inflates false positives at nonzero weight (GitHub issue #45; github.com/broadinstitute/ssGSEA2.0/issues/45). To reproduce this issue, we used a published dataset of feature weights from non-negative matrix factorization (NMF) of a lung adenocarcinoma (LUAD) cohort (Satpathy *et al*. 2025). We randomly constructed 400 gene sets based on the genes detected in this dataset, then ran ssGSEA2.0-orig with weight = 0 or 0.75, as the correlation-vector reordering does not manifest at weight = 0. Both the nominal rate (11.6%) and the Kolmorogorov-Smirnov (KS) test (*p* < 0.0001) at weight = 0.75 demonstrate that ssGSEA2.0-orig is miscalibrated for null gene sets (Figure 1, Table 1). We corrected this behavior by holding the weight vector fixed to its observed order and permuting only gene-set membership (ssGSEA2.0-fixed) and demonstrated its nominal rate (5.2%) is much closer to the theoretical 5% at weight = 0.75. While the KS *p*-value is still significant, the magnitude is greatly reduced, and similar significance at weight = 0 points to an outstanding weight-independent calibration imperfection that nonetheless produces a *p*-value distribution visually indistinguishable from uniform.

**Figure 1.**
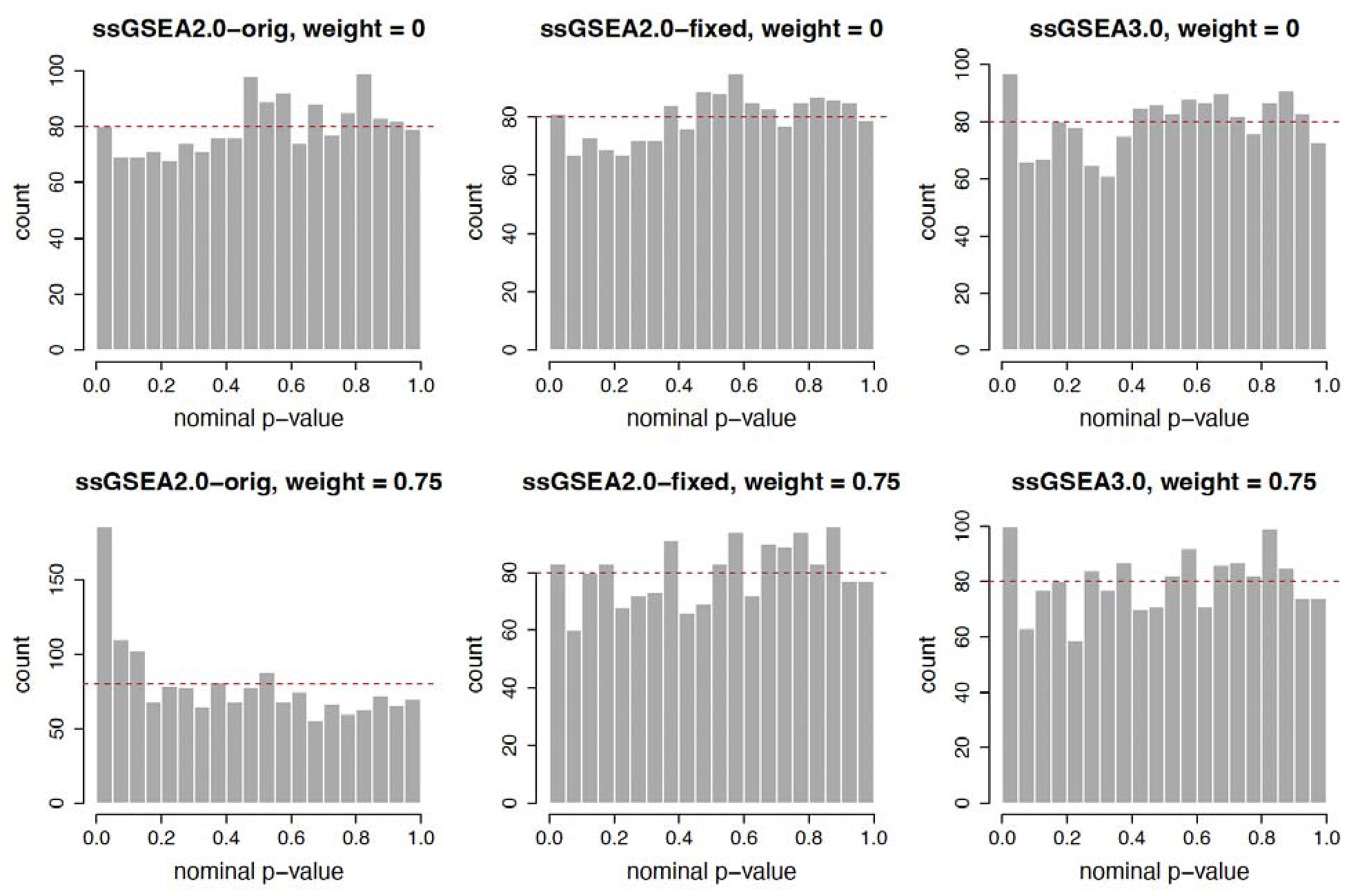
Nominal p-value distributions from null calibration tests by method and weight. The dashed red line marks the expected bin height under a Uniform(0,1) null distribution.

**Table 1.** Null calibration test results. KS: Kolmogorov-Smirnov test.

| Method | Weight | Median *p*-value | KS *D* | KS *p*-value | Nominal<br>*p* < 0.05 rate |
|---|---|---|---|---|---|
| ssGSEA2.0-orig | 0 | 0.5306 | 0.0442 | 0.0039 | 5.00% |
| ssGSEA2.0-orig | 0.75 | 0.4228 | 0.1052 | < 0.0001 | 11.6% |
| ssGSEA2.0-fixed | 0 | 0.5290 | 0.0435 | 0.0047 | 5.06% |
| ssGSEA2.0-fixed | 0.75 | 0.5315 | 0.0394 | 0.0140 | 5.19% |
| ssGSEA3.0 | 0 | 0.5235 | 0.0360 | 0.0318 | 6.00% |
| ssGSEA3.0 | 0.75 | 0.5204 | 0.0217 | 0.4413 | 6.25% |

### The calibration correction in ssGSEA2.0-fixed comes at a detection cost

To determine the effect of the correlation-vector reordering on published biological results, we re-ran the ssGSEA analysis that we previously published on the LUAD NMF dataset (Satpathy *et al*. 2025) using ssGSEA2.0-fixed (Supplementary Table 1). The original analysis was performed against the MSigDB HALLMARK gene sets with weight = 1 at FDR < 0.01. As expected, there was a loss of significant pathways with the correlation-vector reordering fixed (44 in ssGSEA2.0-orig to 36 in ssGSEA2.0-fixed, 18% decrease) (Supplementary Figure 1A). However, the results were highly concordant with pairwise Normalized Enrichment Score (NES) correlation of *r* = 0.98 (Supplementary Figure 1B); further, all 36 significant pathways in ssGSEA2.0-fixed were also significant in ssGSEA2.0-orig (Supplementary Figure 1C). A heatmap of the 28 curated gene sets shown in the original publication is visibly consistent whether using ssGSEA2.0-orig or ssGSEA2.0-fixed (Supplementary Figure 1D). This suggests that the most significant biological signals identified in ssGSEA2.0-orig are not affected by the correlation-vector reordering, and previously published results using ssGSEA2.0-orig are still largely reliable even if a nonzero weight was used.

We additionally performed ssGSEA analysis using weight = 0 as a control, confirming that the correlation-vector reordering does not alter results when weighting is not used (Supplementary Table 1). We noted that, unlike ssGSEA2.0-orig, ssGSEA2.0-fixed returns less significant pathways with a nonzero weight (5% decrease in ssGSEA2.0-fixed versus 16% increase in ssGSEA2.0-orig). While a fraction of significant pathways in ssGSEA2.0-orig are likely false positives due to correlation-vector reordering, we were concerned that the correlation-vector reordering fix may compromise true biological signal, leading to increased false negatives and subsequent loss of statistical power. To further investigate this, we repeated the analysis on the LUAD data using the MSigDB REACTOME collection, which has a higher number of more diverse gene sets, with an FDR < 0.05 (Supplementary Table 2). The pattern we saw in the HALLMARK comparison was even more apparent in REACTOME; despite high NES correlation of *r* = 0.97, there was a 28% decrease in the number of significant pathways between ssGSEA2.0-orig and ssGSEA2.0-fixed at weight = 1 (Figure 2A-B). Further, the gap in significant pathways between weight = 0 and weight = 1 widened for ssGSEA2.0-fixed (10% decrease) compared to HALLMARK (Figure 2A). As before, the majority of pathways significant in ssGSEA2.0-fixed were also significant in ssGSEA2.0-orig (733/734, 99.9%) (Figure 2C). We examined the 286 pathways which were missed by ssGSEA2.0-fixed and found a number of them which were biologically relevant for LUAD, including FGFR1 signaling and cellular senescence (Figure 2D). This suggests that while ssGSEA2.0-fixed does indeed fix the correlation-vector reordering, that fix comes at a detection cost.

**Figure 2.**
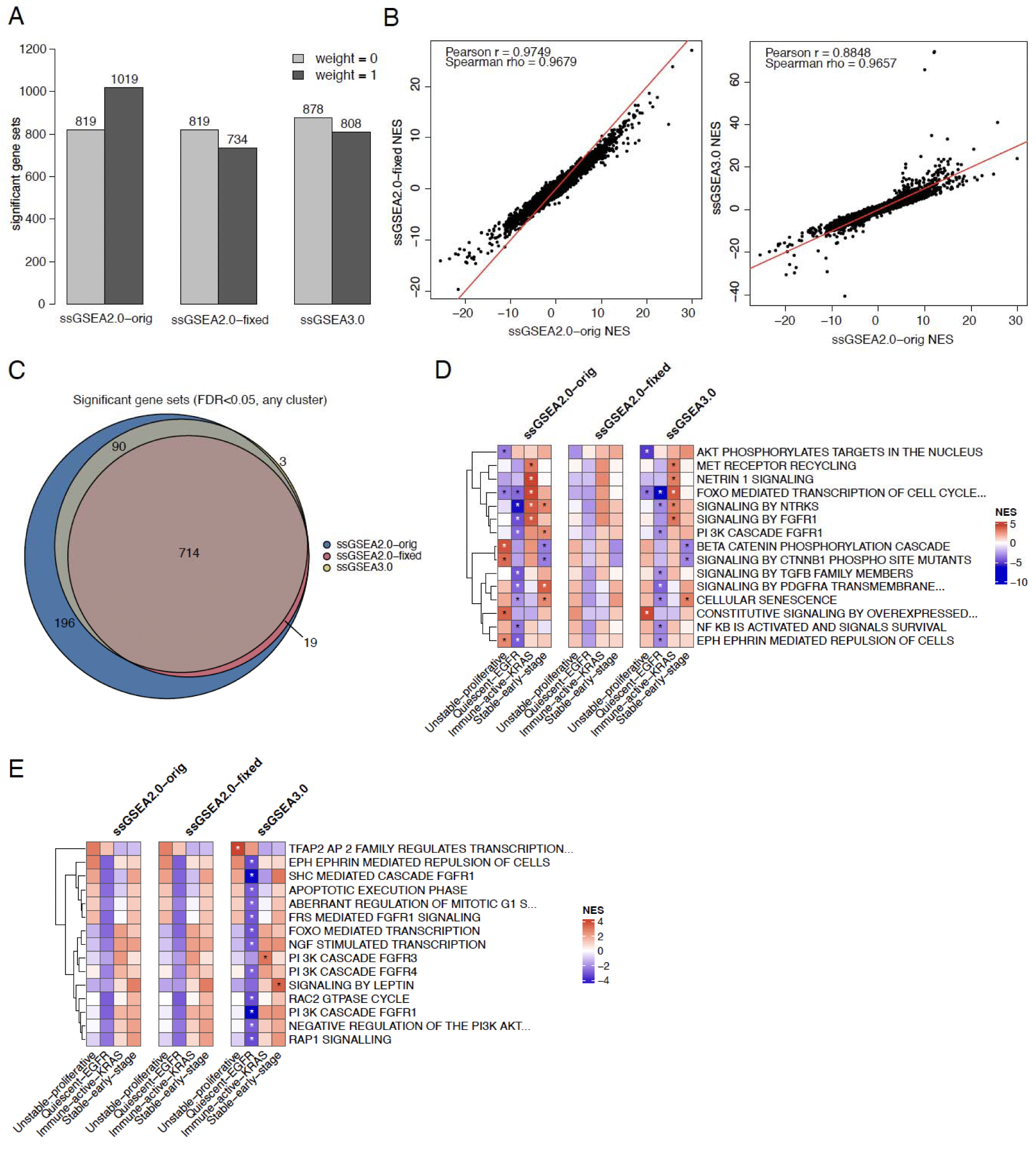
Comparison of the three ssGSEA implementations using the LUAD feature weights from NMF clustering against the MSigDB REACTOME database. (A) Number of significant (FDR < 0.05) gene sets at weight = 0 and weight = 1 for each method. (B) Correlation of Normalized Enrichment Score (NES) across all gene sets and clusters at weight = 1, comparing ssGSEA2.0-fixed (left) and ssGSEA3.0 (right) to ssGSEA2.0-orig. (C) Overlap of significant (FDR < 0.05) gene sets between the three methods. (D) Heatmap of 15 representative gene sets which are significant in ssGSEA2.0-orig and ssGSEA3.0 but missed by ssGSEA2.0-fixed at weight = 1. (E) Heatmap of 15 representative gene sets which are significant in ssGSEA3.0 only at weight = 0. In (D) and (E), * indicates FDR < 0.05.

### Feature-level bootstrap resampling in ssGSEA3.0 restores calibration without sacrificing detection power

Given the observed decrease in detection power of ssGSEA2.0-fixed, we explored whether altering the permutation methodology could restore biologically relevant gene sets while retaining null distribution calibration. One alternative to traditional gene-label permutation is bootstrap resampling of the measured features themselves (Efron and Tibshirani 1994). We thus implemented feature-level bootstrap resampling in place of the existing gene-label permutation algorithm in the new ssGSEA3.0 R package (github.com/broadinstitute/ssGSEA3.0). Bootstrap resampling is more computationally expensive than gene-label permutation as resampling with replacement changes which genes are present in each iteration, requiring us to re-rank the data and re-compute gene-set alignment on every iteration. This results in a ∼1.5 to 2-fold increase in runtime depending on the number and diversity of gene sets (Table 2).

**Table 2.**
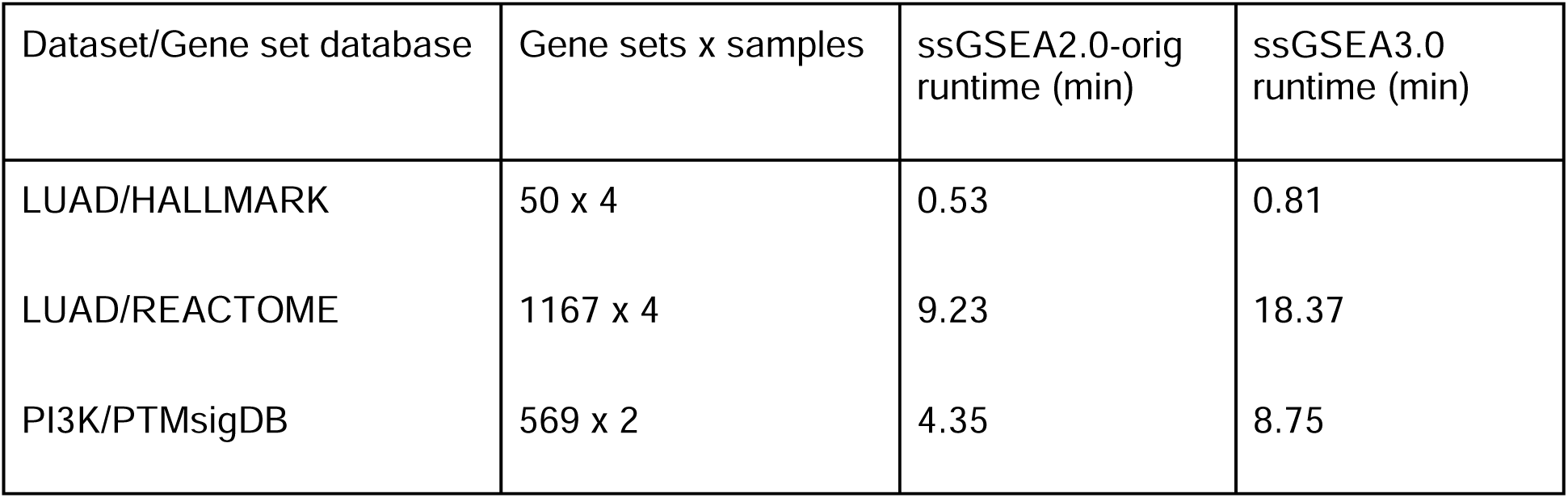
Computational timing benchmarks on gene-label permutation (ssGSEA2.0-orig) and feature-level bootstrap resampling (ssGSEA3.0) algorithms.

We used our simulated null gene sets to demonstrate that ssGSEA3.0 is well-calibrated across weighting schemes, showing a similarly small but nominally significant KS deviation at weight = 0 (*p*-value = 0.0318) that suggests the weight-independent calibration imperfection observed across all three methods is not specific to permutation-based testing (Figure 1, Table 1). However, unlike ssGSEA2.0-fixed, this deviation resolves at weight = 0.75, where ssGSEA3.0’s null p-value distribution is no longer significantly different from uniform (KS *p*-value = 0.4413). ssGSEA3.0 performed on the LUAD NMF dataset against the HALLMARK database with weight = 1 retains 2 additional pathways which were lost with ssGSEA2.0-fixed (Supplementary Figure 1A). While the NES correlation between ssGSEA3.0 and ssGSEA2.0-orig is slightly worse (*r* = 0.95) than for ssGSEA2.0-fixed, the heatmap remains visually consistent (Supplementary Figure 1B, D). The improvements of ssGSEA3.0 over ssGSEA2.0-fixed are most apparent using REACTOME, where ssGSEA3.0 returns 808 significant pathways with weight = 1, a 10% increase over ssGSEA2.0-fixed (Figure 2A). A number of these retained pathways are biologically relevant to LUAD (Figure 2D). Further, ssGSEA3.0 has more significant pathways at weight = 0 than either ssGSEA2.0 method (878 vs 819, 7% increase) which are also biologically relevant, including PI3K signaling and aberrant cell cycle response (Figure 2E). Thus, not only does ssGSEA3.0 fix the null distribution calibration issue present in ssGSEA2.0-orig, but it does so without sacrificing as much detection power as ssGSEA2.0-fixed. Additionally, the results for weight = 0 indicate that the feature-level bootstrapping in ssGSEA3.0 may confer greater detection power than gene-label permutation methods. To test this, we generated a fully synthetic dataset where we embedded a ± 0.5 shift in gene set members (20 members per gene set) and compared detection power in ssGSEA2.0-fixed and ssGSEA3.0. We found that ssGSEA3.0 detected significantly more true positives than ssGSEA2.0-fixed (4,629 vs. 4,211 of 10,000 gene sets; odds ratio = 3.11, McNemar *p* < 0.0001), confirming a genuine power advantage for feature-level bootstrapping over gene-label permutation.

### ssGSEA3.0 confers a power advantage most pronounced for directional PTM-SEA gene sets

Our group previously developed a complementary approach to ssGSEA for Post-Translational Modification (PTM)-centric datasets, such as phosphoproteomics datasets, called PTM-SEA (Krug *et al*. 2019). PTM-SEA uses the same gene-label permutation approach as ssGSEA2.0-orig but performs the enrichment on a PTM site-centric level using the PTMSigDB library of pathways (<u>proteogenomics.shinyapps.io/ptmsigdb/</u>). Importantly, the pathway sets incorporated into PTMSigDB encode directionality of the PTM, so the enrichment score for a particular set is a combination of the enrichment scores computed separately from the “up” and “down” subsets. Thus, we wanted to determine the impact of the feature-level bootstrap resampling in ssGSEA3.0 in the context of PTM-SEA. We used our previously published phosphoproteomics dataset in which T47D breast cancer cells were treated with the PI3Kα inhibitor BYL719 versus DMSO control at 6h and 24h (Krug *et al*. 2019). The input values for PTM-SEA were the log-signed *p*-values from the moderated two-sample *t-*test comparing BYL719 to DMSO at each timepoint. Since we performed PTM-SEA with weight = 0, ssGSEA2.0-orig and ssGSEA2.0-fixed returned equivalent results (differing only by permutation noise, as ssGSEA2.0 cannot be seeded); thus, we used ssGSEA2.0-fixed for the comparison with ssGSEA3.0.

Similar to the ssGSEA results from the LUAD dataset, there was high NES correlation (*r* = 0.91) between the two methods, and the majority of the significant pathways in ssGSEA2.0-fixed were also identified by ssGSEA3.0 (86/93, 92%) (Supplementary Table 3, Figure 3A-B). However, ssGSEA3.0 had a 35% increase in significant pathways over ssGSEA2.0-fixed (Figure 3B); this greatly exceeds the previous 7% increase observed in the LUAD REACTOME weight = 0 comparison. Many of these pathways are biologically relevant and represent multiple components of the PI3K-signaling axis including upstream activators (PDK1/AMPKA2), downstream targets (p70S6K/SGK1), MAPK-pathway crosstalk, and cell-cycle response (Figure 3C). These results suggest that the feature-level bootstrap resampling implemented in ssGSEA3.0 confers a power advantage for directional gene sets.

**Figure 3.**
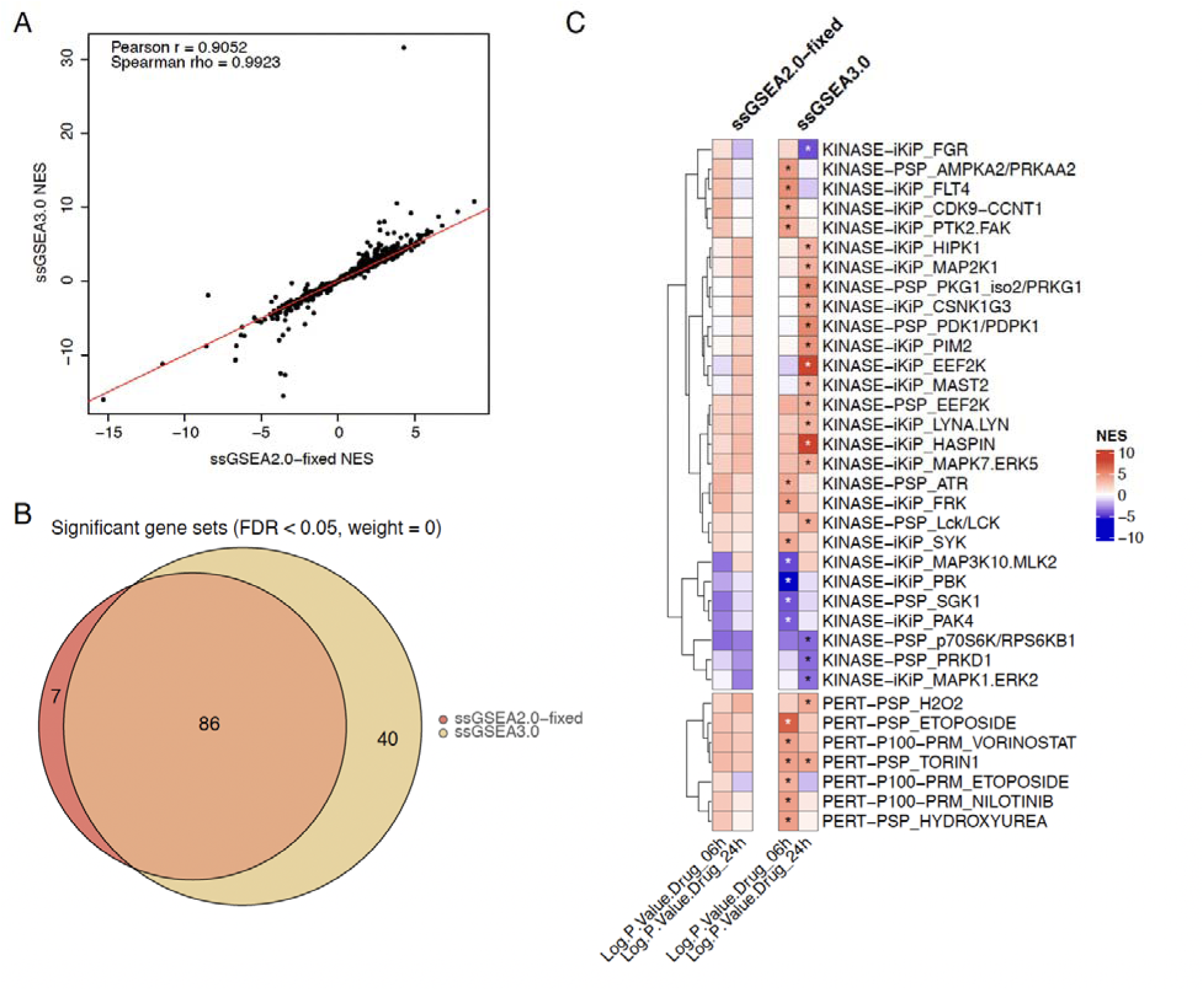
Comparison of PTM-SEA performed using ssGSEA2.0-fixed or ssGSEA3.0 on the PI3Kα-inhibitor phosphoproteomics dataset from Krug *et al* (2019) with weight = 0. (A) Correlation of Normalized Enrichment Score (NES) across all gene sets between ssGSEA2.0-fixed and ssGSEA3.0. (B) Overlap of significant (FDR < 0.05) gene sets between the two methods. (C) Heatmap of 35 representative gene sets which are significant in ssGSEA3.0 but not ssGSEA2.0-fixed. * indicates FDR < 0.05.

The additional detection power in ssGSEA3.0 could reflect either a bootstrap miscalibration (an inflated false-positive rate) or a genuine power advantage. To address the first concern, we generated a fully synthetic dataset and performed a null-calibration simulation at two gene set sizes (20 or 100), both directional and non-directional, for either gene-label permutation or feature-level bootstrap resampling. All configurations were well-calibrated for null gene sets whether permutation or bootstrap approaches were used (Table 3). To test if there is a power advantage and if it is specific to directional scoring, we again used our simulated, shifted gene-sets but incorporated both directional and non-directional gene sets. The feature-level bootstrap resampling returned significantly more pathways than the gene-level permutation approach, regardless of the directionality (Table 4). Thus, we conclude that the feature-level bootstrap resampling implemented in ssGSEA3.0 confers a power advantage over the traditional gene-label permutation approach. Further, on real biological data, this advantage is more pronounced for directional gene sets such as those used in PTM-SEA.

**Table 3.**
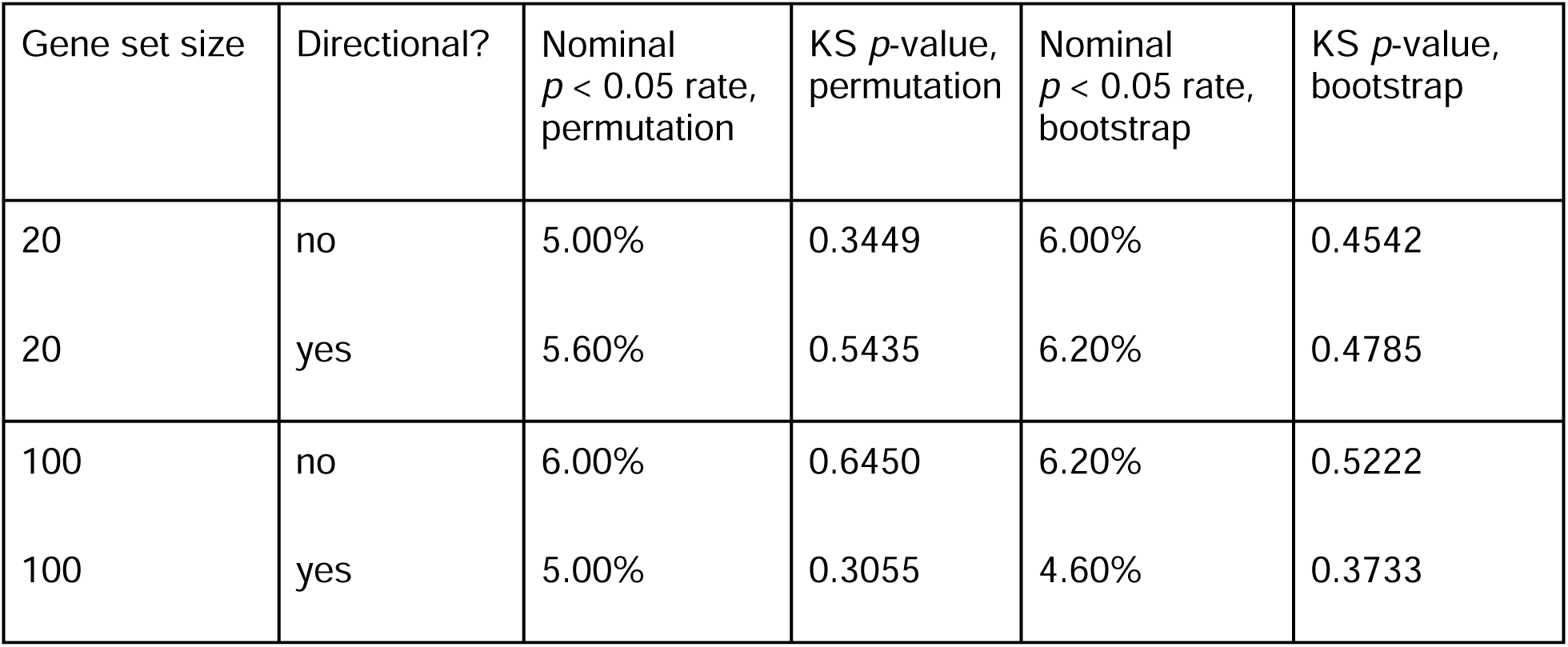
Null calibration test results using both permutation and bootstrap approaches on simulated non-directional or directional gene sets of different sizes with weight = 0. KS: Kolmogorov-Smirnov test.

**Table 4.** Power detection test using a known embedded ± 0.5 signal with both permutation and bootstrap approaches on simulated non-directional or directional gene sets with weight = 0.

| Gene set size | Directional? | Number of gene sets significant for bootstrap only | Number of gene sets significant for permutation only | Odds ratio | McNemar *p*-value |
|---|---|---|---|---|---|
| 20 | no | 428 | 229 | 1.87 | < 0.0001 |
| 20 | yes | 390 | 296 | 1.32 | 0.0004 |

## Discussion

We demonstrate here that the current ssGSEA2.0 implementation (ssGSEA2.0-orig) is miscalibrated under permutation testing for non-zero weights. In this implementation, reordering the weight vector along with the feature ranks results in an overly narrow null distribution and inflates false positives. Holding the weight vector fixed is a straightforward fix (ssGSEA2.0-fixed) that restores calibration but reduces detection power, particularly on larger, more diverse gene set collections. ssGSEA3.0’s feature-level bootstrap resampling resolves both problems simultaneously: it is well-calibrated and retains more significant gene sets than ssGSEA2.0-fixed. Further, we show that feature-level bootstrapping confers a power advantage over the traditional gene label permutation approach even at weight = 0. Given these results, we recommend ssGSEA3.0 as the default implementation moving forward, particularly for analyses incorporating non-zero weights or directional gene set collections such as PTMSigDB. We provide ssGSEA3.0 as a direct replacement for existing ssGSEA2.0 workflows, accepting the same GCT input format and producing the same GCT output files. It is available as an open-source R package with a worked vignette reproducing the HALLMARK, REACTOME, and PTM-SEA comparisons presented here.

These findings carry a practical implication for the substantial body of published work that has used ssGSEA2.0-orig with weight > 0, which is the standard setting. The high concordance we observe between ssGSEA2.0-orig and ssGSEA2.0-fixed on real data suggests that the majority of previously reported significant pathways reflect real biological signals rather than an artifact of the reordering miscalibration. We would not expect this finding to overturn prior conclusions built on strong, well-replicated enrichment signals; however, we recommend cautious interpretation of results near the significance threshold.

Some limitations qualify these findings. While we sought a rigorous cross-comparison by evaluating all methods against each other with both synthetic and real biological data, we were unable to extend our synthetic ground-truth approach to directly compare ssGSEA2.0-orig against ssGSEA2.0-fixed or ssGSEA3.0. This is because the correlation-vector reordering in ssGSEA2.0-orig alters the observed enrichment statistic itself at a non-zero weight, not only its null distribution. Therefore no choice of significance threshold can separate a true power difference from this altered scoring. The detection-cost comparisons in this manuscript involving ssGSEA2.0-orig therefore rest on real-data concordance alone, which we consider sufficient to support our conclusions but which a controlled synthetic experiment could not independently confirm. Additionally, ssGSEA3.0 is not perfectly calibrated. Nominal rates were slightly above 5% (6.0–6.3%), and the KS test showed a small deviation at weight = 0. A similar deviation appeared in both ssGSEA2.0 variants, so it is not specific to bootstrapping, but a modest excess of false positives cannot be excluded. Finally, in the biological comparisons, relevance of the significant pathways was judged by manual inspection rather than against an independent ground truth. ssGSEA3.0 also incurs a modest increase in computational cost, running about 1.5- to 2-fold slower than ssGSEA2.0-orig.

We hypothesize the power difference between bootstrap resampling and permutation testing reflects how each method builds its null distribution. Standard GSEA tests significance by shuffling sample labels across a cohort and recomputing the gene ranking each time, which is only reliable when multiple samples are available. Since ssGSEA scores one sample at a time, there are no sample labels to shuffle; instead, ssGSEA2.0 builds its null by randomly reordering the ranked gene list itself, which appears to be a weaker substitute given the drop in detection power of ssGSEA2.0-fixed. Bootstrap resampling avoids the problem by resampling the feature values directly to capture the variability of the score, leading to a more accurate null.

The power advantage we observe with bootstrap resampling is substantially larger for directional PTM-SEA scoring (+35%) than for non-directional REACTOME scoring (+7%), even though our synthetic simulation shows the underlying bootstrap advantage is not inherently directional-specific. One possible explanation is that directional scoring’s combination of separately computed “up” and “down” sub-scores compounds the power gained from a more accurate null at each step, but we did not test this explicitly and leave it as a future direction.

More broadly, other single-sample enrichment methods that rely on gene-label permutation for significance testing may share the same calibration and power trade-offs we characterize here, and feature-level resampling may offer a general alternative worth evaluating in those settings. Extending this validation to additional data types and cohorts would further establish the generality of these findings.

## Data availability statement

The LUAD NMF weight matrix used here corresponds to the ssGSEA results shown in Figure 1 of (Satpathy *et al*. 2025), derived via the PANOPLY pipeline (Mani *et al*. 2021) from proteogenomic data publicly available through LinkedOmics (https://www.linkedomics.org/data_download/CPTAC-ICPC-LUAD/). This derived matrix was not deposited as a supplementary dataset in the original publication; it can be accessed via the ssGSEA3.0 software package, archived at Zenodo under DOI: 10.5281/zenodo.23734262 (github.com/broadinstitute/ssGSEA3.0). The PI3Kα-inhibitor phosphoproteomics dataset is available as Supplementary Table 6 of (Krug *et al*. 2019) and is additionally included in the ssGSEA3.0 package.

## Acknowledgements

Claude Code was used to evaluate the codebase for bugs and optimizations, re-structure the codebase as an R package, write unit tests, and format documentation. The authors have reviewed these additions and changes for accuracy.

We thank Greg Sepich-Poore and Tyler Sagendorf for identifying and reporting the weight-dependent calibration issue in ssGSEA2.0 (GitHub issue #45, github.com/broadinstitute/ssGSEA2.0/issues/45), which motivated this work.

This work was partially supported by the National Institutes of Health [U24-CA271075 to D. R. M.].

## Notes

https://doi.org/10.5281/zenodo.23734262

https://doi.org/10.5281/zenodo.23734330

https://github.com/broadinstitute/ssGSEA3.0

https://www.linkedomics.org/data_download/CPTAC-ICPC-LUAD/

https://doi.org/10.1074/mcp.TIR118.000943

### Competing Interest Statement

The authors have declared no competing interest.

