## Supplementary Data for "ssGSEA3.0: Feature-Level Bootstrap Resampling for Robust Assessment of Statistical Significance"

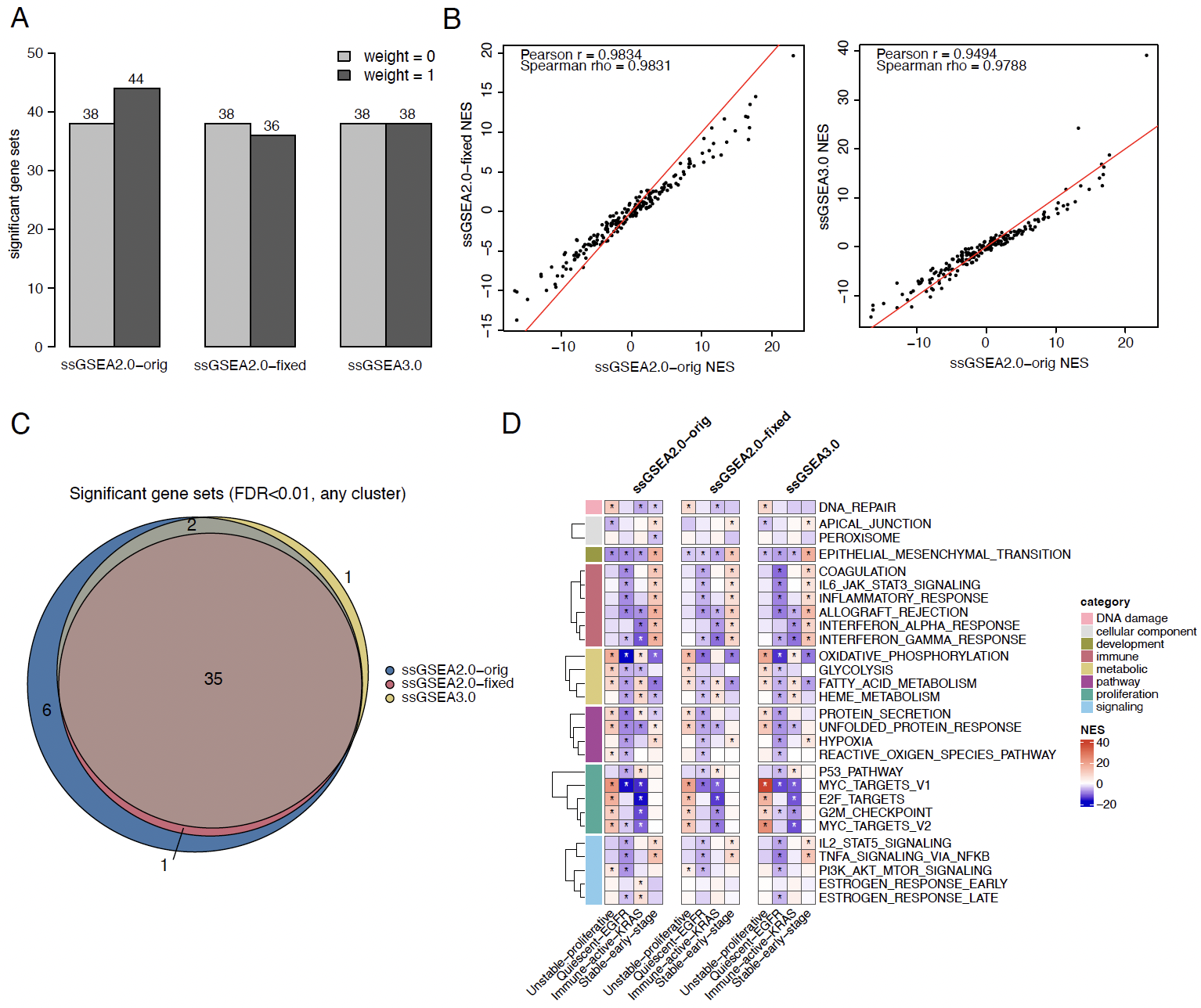

**Supplementary Figure 1.** Comparison of the three ssGSEA implementations using the LUAD feature weights from NMF clustering against the MSigDB HALLMARK database. (A) Number of significant (FDR < 0.01) gene sets at weight = 0 and weight = 1 for each method. (B) Correlation of Normalized Enrichment Score (NES) across all gene sets and clusters at weight = 1, comparing ssGSEA2.0-fixed (left) and ssGSEA3.0 (right) to ssGSEA2.0-orig. (C) Overlap of significant (FDR < 0.01) gene sets between the three methods. (D) Heatmap of the 28 gene sets shown in Satpathy *et al.* (2025) grouped by term category. * indicates FDR < 0.01.

**Supplementary Table 1.** Results from ssGSEA performed on the LUAD NMF dataset against MSigDB HALLMARK.

**Supplementary Table 2.** Results from ssGSEA performed on the LUAD NMF dataset against MSigDB REACTOME.

**Supplementary Table 3.** Results from PTM-SEA performed on the PI3Kα-inhibitor dataset.
