## Supplementary Figure 1 for "ssGSEA3.0: Feature-Level Bootstrap Resampling for Robust Assessment of Statistical Significance"

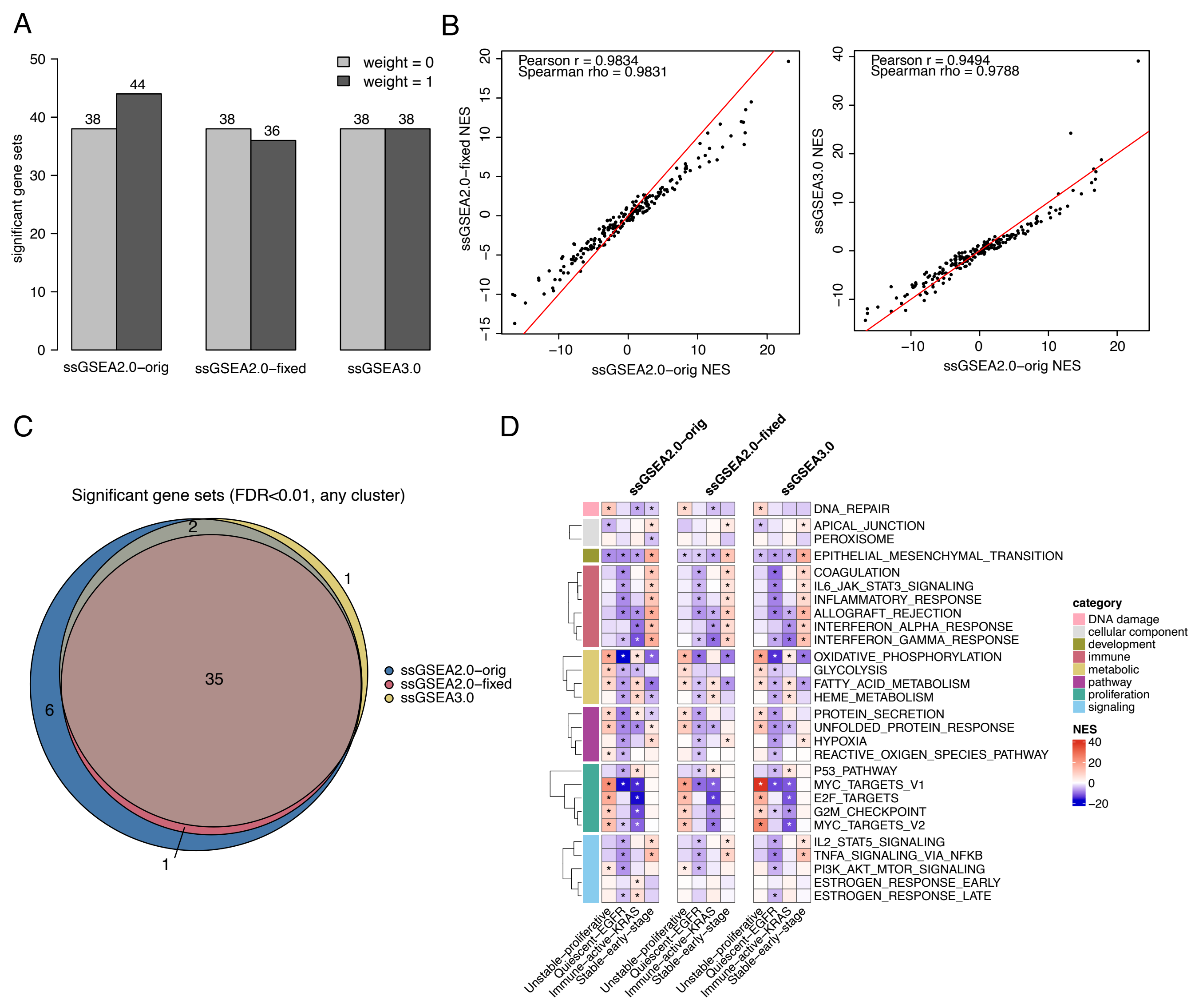
A
weight = 0
weight = 1
significant gene sets
ssGSEA2.0–orig
ssGSEA2.0–fixed
ssGSEA3.0
38
44
38
36
38
38
B
Pearson r = 0.9834
Spearman rho = 0.9831
ssGSEA2.0–fixed NES
ssGSEA2.0–orig NES
Pearson r = 0.9494
Spearman rho = 0.9788
ssGSEA3.0 NES
ssGSEA2.0–orig NES
C
Significant gene sets (FDR<0.01, any cluster)
2
1
35
6
1
ssGSEA2.0–orig
ssGSEA2.0–fixed
ssGSEA3.0
D
ssGSEA2.0–orig
ssGSEA2.0–fixed
ssGSEA3.0
DNA_REPAIR
APICAL_JUNCTION
PEROXISOME
EPITHELIAL_MESENCHYMAL_TRANSITION
COAGULATION
IL6_JAK_STAT3_SIGNALING
INFLAMMATORY_RESPONSE
ALLOGRAFT_REJECTION
INTERFERON_ALPHA_RESPONSE
INTERFERON_GAMMA_RESPONSE
OXIDATIVE_PHOSPHORYLATION
GLYCOLYSIS
FATTY_ACID_METABOLISM
HEME_METABOLISM
PROTEIN_SECRETION
UNFOLDED_PROTEIN_RESPONSE
HYPOXIA
REACTIVE_OXIGEN_SPECIES_PATHWAY
P53_PATHWAY
MYC_TARGETS_V1
E2F_TARGETS
G2M_CHECKPOINT
MYC_TARGETS_V2
IL2_STAT5_SIGNALING
TNFA_SIGNALING_VIA_NFKB
PI3K_AKT_MTOR_SIGNALING
ESTROGEN_RESPONSE_EARLY
ESTROGEN_RESPONSE_LATE
category
DNA damage
cellular component
development
immune
metabolic
pathway
proliferation
signaling
NES
40
20
0
−20
Unstable–proliferative
Quiescent–EGFR
Immune–active–KRAS
Stable–early–stage
